# BrainProt(™) 3.0: Understanding Human Brain Diseases using comprehensively curated & Integrated OMICS datasets

**DOI:** 10.1101/2023.06.21.545851

**Authors:** Deeptarup Biswas, Sanjyot Vinayak Shenoy, Aparna Chauhan, Ankit Halder, Biplab Ghosh, Advait Padhye, Shreeman Auromahima, Deeksha Yadav, Souvik Sasmal, Sampurna Dutta, Neha Kumari, Hiren Bhavaskar, Ayan Prasad Mukherjee, Tunuguntla Rishi Kumar, Sanjeeva Srivastava

## Abstract

BrainProt 3.0 is an integrative and simplified omics-based knowledge base of the human brain and its associated diseases. The current version of BrainProt includes six domains, which provide simplified, robust, and comprehensive data visualization to understand the human brain and its diseases/disorders based on proteomics, transcriptomics, public data curation, and integration strategies. Firstly, the HBDA (Human Brain Disease Atlas), index and navigator of BrainProt provides a resource table for 56 brain diseases. Secondly, Brain Disease Marker Curator (BDMC) and Brain Disease Drug Finder (BDDF) include a total of 20,202 diseases associated genes, more than 1,30,000 Chemical Target interactions, and around 2,145 Clinical Trial Information for more than 50 Brain Diseases. Thirdly, Brain Disease Transcriptome Map (BDTM) and Brain Disease Proteome Map (BDPM) integrate multi-omics data for 11 and 6 alarming brain diseases respectively. Currently, these two domains feature an expressional profile of 52 datasets, information of 1,868 samples, 3,657 DEPs, and 6,256 DEGs. Lastly, BrainProt also modifies and integrates the proteome and phosphoproteome data of the Inter-hemispheric Brain Proteome Map (IBPM). Overall, BrainProt is the first knowledgebase that connects different omics level information of brain diseases and provides a powerful scoring-based ranking platform to select and identify brain disease-associated markers, along with exploration of clinical trials, and drugs/chemical compounds to accelerate the identification of new disease markers and novel therapeutic strategies. The objectives of BrainProt are to support and follow the footsteps of the HBPP (Human Brain Proteome Project) by integrating different datasets to unravel the complexity of Human Brain and its associated diseases.

## INTRODUCTION

The human Brain and its associated functionalities have always been a major hub of attraction for research among scientists worldwide. The evolution of the human brain and its function has always perplexed the scientific world.. The evolution of Omics-based technologies and the revolution of data analysis strategies have fostered the generation and understanding of big data from the perspective of clinical biology (1). In context of the human brain diseases, Human brain tumors (BT) and neurodegenerative disorders (NDD) have become the major concerns of the medical fraternity and require a global effort to deal with them (2, 3). The foundation of HBPP under the leadership of HUPO has expedited the identification of new biomarkers and novel therapeutic strategies focusing primarily on omics technologies (4). The analysis and interpretation of these brain-related disease datasets have shed light on comprehending the molecular level alteration to identify better diagnostic strategies and unveil novel therapeutic modalities. However, a lot more information is concealed in this humongous data, which is available in different data repositories, and manuscripts could be revisited and reanalysed on a global level to retrieve more information and provides more confidence needed to pin-point out a marker for a certain disease (5).

In the era of integrative omics and big data analysis, a need for robust, fast, and interpretable omics-based visualization portals to decipher the variation of the expressional profile of different brain tumors and disorders are of utmost importance. Many research groups around the globe have carried out the initiative toward the same path, but the majority have focused either on Glioma/Glioblastoma, being the most aggressive form of a brain tumor, or on a single disease like Alzheimer’s Disease or Parkinson’s disease. Some recent and popular OMICS-driven databases like GliomaDB (6), GlioMarker (7), Glioma-BioDP (8), Ivy Glioblastoma Atlas Project (9), GlioVis (10), and the pathology atlas of Human Protein Atlas (HPA) (11) have integrated Glioma and GBM information. On the other hand, web portals like BCGene (12) which has integrated 20 brain cancer gene-associated information, have focused on data curation strategies only. Brainbase despite being one of the most comprehensive, manually curated knowledgebase of a large number of brain diseases, but still limited to Glioma only in regards to omics expressional data like Genome, Transcriptome, and, Epigenome (13). The necessity of an integrated omics-driven database for guiding the neuroscience community to understand, compare and correlate dreadful brain diseases is required to substantiate different reported markers and expedite clinical research.

Here, we have updated the BrainProt knowledgebase by developing and integrating five new domains and modifies IBPM domain into a user-friendly and robust web interface. Currently, version 3.0 of BrainProt integrates transcriptome and proteome data mined and analyzed from Public repositories in context to diseases like Alzheimer’s disease, Parkinson’s disease, Multiple sclerosis, Huntington’s disease, Amyotrophic lateral sclerosis, Frontotemporal lobar degeneration, Meningioma, Medulloblastoma, Glioma, Glioblastoma and Pituitary adenoma. Additionally, popular disease-gene-associated mining platforms and drug databases were used to curate disease markers and drugs closely related to the diseases. Furthermore, this web portal also provides clinical trial information on brain disease searches along with Trial Phase. Finally, all these data have been comprehensively integrated into Brainprot (http://www.brainprot.org/) with an aim to quench the thirst of the scientific community’s interest in understanding brain disease biology.

## MATERIALS AND METHODS

### Development of HBDA and Modification of IBPM data visualization

A list of 56 human brain diseases was curated from Medical Subject Headings (MeSH) available in Table 1. Each of the diseases has been integrated with a brief summary along with a list of 10 databases hyperlinked under a resource table. The IBPM section of BrainProt has been developed and released and published previously (14, 15), which has been modified and updated with interpretable visualization plot, simplified data tables and easy link to 10 knowledgebases under Protein Resource Hub in version 3.0.

### Database curation, dataset mining, and filtration for BDTM and BDPM

A list of 11 alarming human brain diseases has been selected from published manuscripts and disease databases which includes brain tumors and neurodegenerative diseases like Alzheimer’s disease, Parkinson’s disease, Multiple sclerosis, Huntington’s diseases, Amyotrophic lateral sclerosis, Fronto-temporal lobar degeneration, Meningioma, Medulloblastoma, Glioma and Glioblastoma and Pituitary adenoma.

The data mining of proteomics and transcriptomics datasets for the BDTM and BDPM was performed using multiple public data repositories, like ProteomeXchange Consortium (16), PRIDE (17), ArrayExpress (18), Gene Expression Omnibus (GEO) (19), and Omics Discovery Index (OmicsDI) (20). The keyword used for the disease search was “Disease Name.” Datasets were primarily selected keeping the following criteria i) species: *Homo sapiens*, ii) sample type: tissue, and iii) proper clinical meta-data information along with the manuscript. The sample type “tissue” includes both Fresh Frozen (FF) and Formalin-fixed, paraffin-embedded (FFPE). Datasets that fail to provide transparent information about the datasets in the manuscript or sample meta-data file couldn’t be taken forward for the analysis. Unlike BDTM, BDPM includes 6 major dreadful brain diseases which includes Alzheimer’s disease, Parkinson’s disease, Amyotrophic lateral sclerosis, Meningioma, Glioma/Glioblastoma and Medulloblastoma based on proteomics data availability.

### Data Processing and Analysis of omics data for the development of BDTM and BDPM

The current version of BDTM includes transcriptome data analyzed using a uniform data pre-processing, quality check, and statistical analysis pipeline for all the datasets. In brief, the raw CEL files from different databases (NCBI GEO, arrayExpress, Omics DI) were downloaded, extracted, and then loaded into the R. The quality check of the arrays was done using arrayQualityMetrics (21). The Affy package was used for data pre-processing, and the RMA method was used for normalization. Differentially expressed genes were identified by performing moderated t-test using the limma package integrated with AffylmGUI (22). The datasets which couldn’t be analyzed using our pipeline, specifically some exon arrays, were excluded from the study. The datasets containing more than one Affymetrix platform were analyzed independently, given that each platform had matched disease and control samples.

The Proteomics datasets for BDPM were re-analyzed in the MaxQuant (Version:1.6.12) to minimize the cross-search engine variability (23). The Raw files were processed within Label-Free-Quantification (LFQ) parameters, setting label type as “standard” with a multiplicity of 1. The match between runs was selected. Trypsin was used for digestion with a maximum missed cleavage of 2. Carbamidomethylation of cysteine (+57.021464 Da) was set as the fixed modification, whereas oxidation of methionine (+15.994915 Da) was set as the variable modification. The False-Discovery-Rate (FDR) was set to 1 % for the proteins, PSM, and site decoy fraction to ensure high protein identification/quantification reliability. Decoy mode was set to “reverse”, and the minimum peptide length was kept at 7AA.

The significance annotation for BDTM data has been provided using the moderated T-test to calculate pvalue whereas Welch’s T-test has been used to calculate the pValue for BDPM datasets. The differential expression has been studied according to the diiference in expression in disease samples against the control samples. The meta-data of all the datasets are formatted to prepare a consolidate sample information available under each study IDs which are hyperlinked to GEO database for BDTM and ProteomeXchange for BDPM. The Pvalue annotation mapping states *** = p-value < 0.001; ** = p-value < 0.01; * = p-value < 0.05; ns = p-value > 0.05.

### Development of Brain Disease Marker Curator (BDMC)

Brain Disease Marker Curator integrates the power of popular knowledgebases and marker identifier algorithms like Harmonizome (24), BIONDA (25), Pubpular (26), DisGeNET (27), Disease2.0 (28), Comparative Toxicogenomics Database (CTD) (29) and eDGAR (30). The score of each of these databases has been normalized to prepare a consolidated sheet for each of the diseases. The genes present in the eDGAR database and BIONDA database are considered as Disease markers. In addition, the genes of CTD with a (M) marker/mechanism and/or (T) therapeutic annotations under the Direct Evidence column are considered as CTD marker. However, the Disease2.0 markers are selected if the gene of a particular disease has a confidence 3 or more than 3 stars. The BDMC score has been calculated considering the scaled scores and frequency of all 7 databases. The visualization of BDMC domain has been portrayed in the form of user defined axis for scatter plots along with bar plot of top 10 markers for each disease

### Development of Brain Disease Drug Finder (BDDF)

The “Disease Therapeutic Compound” section of the domain has been developed with manual curation and integrates reviewed data. The top markers and therapeutic targets from BDMC and published manuscripts have been taken forward to identify the drugs and chemical compounds available. The manual curation of all the chemical agents reported against the targets are mined from the databases like Ch EMBL (31), Stitch (32), TTD (33), and BindingDB (34). BDDF aimed to provide the toxicity status and the mechanism of action of those reported agents from OpenTarget (35) and CLUE (36). The scores implies the pIC50 score (9-log10(Ic50)) from ChEMBL,BindingDB,and TTD with a larger score beow 9 implying stronger association between the compound and the target and the score from stitch has been scaled between 0 to 1 with a higher score suggesting for better association. The reviewed consolidated sheet has been prepared for 11 alarming brain diseases which includes Alzheimer’s Disease, Amyotrophic Lateral Sclerosis, Frontotemporal Dementia, Glioblastoma, Glioma, Huntington’s Disease, Medulloblastoma, Meningioma, Multiple Sclerosis, Parkinson’s Disease and Pituitary Adenoma.

The “Clinical Trial Information” section has been developed for 53 diseases by integrating the information from Ch EMBL (31) and clinicaltrial.gov (https://clinicaltrials.gov/) to generate an integrated consolidate dsheet. The common studies available in these two databases has only been considered The CHEMBL ID and NCT ID have been hyperlinked for easy access of more information for any studies. 3 out of the 56 Diseases of HBDA has been removed for lack of Clinical Trial information. The BDDF search bar requires an input of protein/gene available in Human database as a query which provides with a series of information and scores like pIC50, Stitch Scores, Mechanism of Action, Clinical Phases from ChEMBL(31), BindingDB(32), Stitch(33), TTD (34) and CLUE (36) database. The BDDF has been developed with a hybrid data accessibility model where few section fetches data directly from the primary database to maintain updated information; however, most of the data are downloaded, curated and reviewed from manuscript and other data repositories.

### Database construction and web interface implementation

BrainProt is based on Django V4.0 through the Apache 2 Web Server (https://tomcat.apache.org/) as the backend. The web interface is a dynamic single-page application based on React (V18.2), while BootStrap V5.0 is used as the base for styling using reactstrap. The interactive data visualization plots like bar plots, lollipop plot, pie chart and colour maps has been provided via AirBnB’s Visx, a node package for React.

## DATABASE CONTENTS AND USAGE

### Development of BrainProt domains

The current version of BrainProt includes five new domains with the existing Inter-Hemisphere Brain Proteome Map (IBPM) (14, 15), which deciphers the expression profile of proteins and phosphosites identified in the left and right hemispheres of different neuroanatomical regions and sub-regions of the Human Brain (Figure 1). The five new domain includes:

- Human Brain Disease Atlas (HBDA): This domain is the index and navigator of BrainProt which includes 56 brain diseases along with disease resource table.
- Brain Disease Marker Curator (BDMC): The domain allows users to identify and select markers of 56 brain diseases based on ranking strategies, stimulating biomarker discovery and therapeutic target identification.
- Brain Disease Transcriptome Map (BDTM): This domain of BrainProt shows the gene expression profile of 11 Brain tumors and Neurodegenerative disorders mined d from ArrayExpress and Gene Expression Omnibus (GEO) and analysed independently.
- Brain Disease Proteome Map (BDPM): This domain shows the protein level expression profile of 6 alarming Brain tumors and Neurodegenerative disorders mined from ProteomeXchange, PRIDE, and OmicsDi and followed by analysis in a particular and independent manner.
- Brain Disease Drug Finder (BDDF): The domain provides clinical trial information on more than 50 brain diseases and allows users to curate, select and identify probable drug and chemical compounds against top therapeutic targets of 11 alarming brain diseases.Also, the known compounds/chemical agents against a particular target can be identified through search of a particular protein/gene

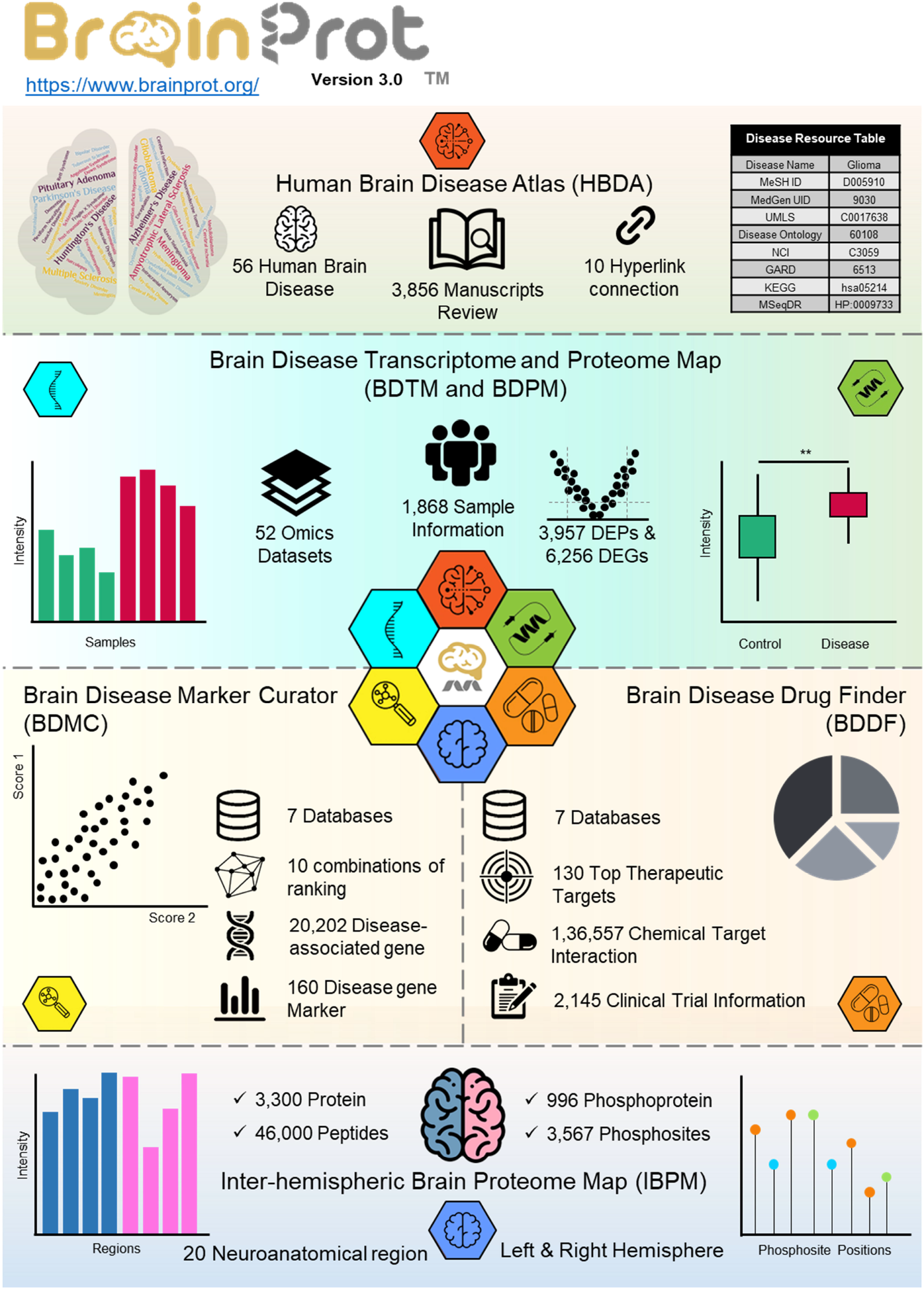

### Selection, Identification and Visualization of human brain disease associated markers using score based ranking strategy

Human Brain Disease Atlas (HBDA) integrates a list of 56 human brain diseases manually curated from Medical Subject Headings (MeSH) based on its classes. The domain has also computed a list of 10 different unique disease IDs dispersed in multiple sources which includes MedGen UID, UMLS, Disease Ontology (DO), NCI, GARD, KEGG, MSeqDR, HP, Monarch Initiative, OMIM along with MeSH ID. This resource table can redirect to different associated information of the selected disease available. The identification of disease-associated genes is one of the most critical steps toward biomarker discovery. Brain Disease Marker Curator (BDMC) integrates information of 7 databases and 20,202 Disease-associated genes to develop a powerful platform for disease marker identification. The section provides an option of 10 different combinations and a user-defined gene numbers to select and visualize disease associated marker through scatter plot; however, the bar plot represents the top 10 marker genes of a particular disease. The calculated BDMC score and frequency of each gene can be used to sort and select top markers for a disease. A total of 160 markers were found from 56 diseases which had a BDMC score more than 7.5.

### Transcriptome and Proteome Profile of alarming human brain diseases

BDTM and BDPM, the major domains of BrainProt, aim to provide a holistic view of inter-disease and intra-disease expressional profile comparison of gene/proteins in the disease and the healthy state Currently, the domains include a total of 52 omics datasets (transcriptomics and proteomics) from 11 daunting brain diseases for BDTM and 6 brain diseases for BDPM. These two domains also integrate the meta-data information of 1,868 sample which can be used for better clinical correlation and interpretation. The domains allow users to visualize the analysed, processed intensity in the form of bar plots along with group-wise box plots to compare the expressional difference between Disease and Control. A list of 6,256 significant genes and 3,957 significant proteins were identified under BDTM and BDPM respectively. The Expression of TP53, IDH1, AKT1 and EGFR, known markers in cancer, has been found to be upregulated primarily in prevalent brain tumors like Glioblastoma, Glioma, and Meningioma under BDTM. The literature-curated markers of Neurodegenerative diseases have also been explored primarily in Alzheimer’s disease and Parkinson’s Disease to compare the expressional profile (37, 38). The expressional difference between the disease and control group of Parkinson’s Disease markers like Tyrosine 3-monooxygenase (TH), Netrin-1 (NTN1), Netrin receptor DCC and Cytoplasmic protein NCK2, were also found to be correlating with the known literature. The known communicated markers of Alzheimer’s Disease, like Amyloid-beta precursor protein (APP), Aquaporin-1 (AQP1), Dystrobrevin alpha (DTNA), Calcium/calmodulin-dependent protein kinase kinase 2 (CAMKK2), and Calmodulin regulator protein PCP4, also matched with most of the datasets with their reported expression level.

### Exploration of Clinical Trial Information, Drug and Chemical Compound of human brain diseases

Brain Disease Drug Finder (BDDF) includes information of 7 databases and comprises of three different sections, which includes exploration of Clinical Trial Information of more than 50 human brain diseases, Selection of drug/chemical compound against the top curated markers of 11 concerning brain diseases and identification of chemical compounds against gene/protein of interest using BDDF search bar. These three sections together make BDDF a dynamic and comprehensive platform for exploring therapeutic aspect of human brain diseases. A total of 2,145 Clinical Trial Information from 53 brain diseases has been integrated under this section which portrays the trial phases and status information through bar plot, pie chart and consolidated tables. In addition, the disease therapeutic compound section includes more than 1,30, 000 chemical compounds and drugs against 130 therapeutic candidates. It shall help the researchers to explore the options of inhibiting a particular known marker with the known chemical agents blended under a single head.

### Role of BrainProt V3.0 toward a better understanding of Human Brain Disease

The combination and power of all the domains together could support and aid neuroscientists, researchers, and clinicians in reaching to the hypothesis and comprehending brain disease rapidly. Tumor protein p53 (TP53) and Epidermal growth factor receptor (EGFR), a known and widely studies marker for brain tumor, have been taken forward to understand the application and usage of BrainProt v3.0 domains (39–41). TP53 and EGFR has been found among the top 10 marker in BDMC for Glioma with a BDMC score of 9.125 and 9.114 respectively. Furthermore, the expression profile of TP53 in BDTM was found to be upregulated in Glioma, Glioblastoma and Meningioma Datasets with a pValue less than 0.05. The expression of EGFR has also been found to be upregulated in BDPM for Glioma, Glioblastoma and Meningioma datasets. The Clinical Trial information recorded a total of 186 study for Glioma, which shows 1 competed Phase 4 trial (NCT00283543), the investigation of DTI-015 (CARMUSTINE). In addition, the total number of chemical compound/Drugs found against TP53 for Glioma are 559 and EGFR for Glioblastoma are 12189 under the Disease Therapeutic Compounds section of BDDF (Figure 2A and B).

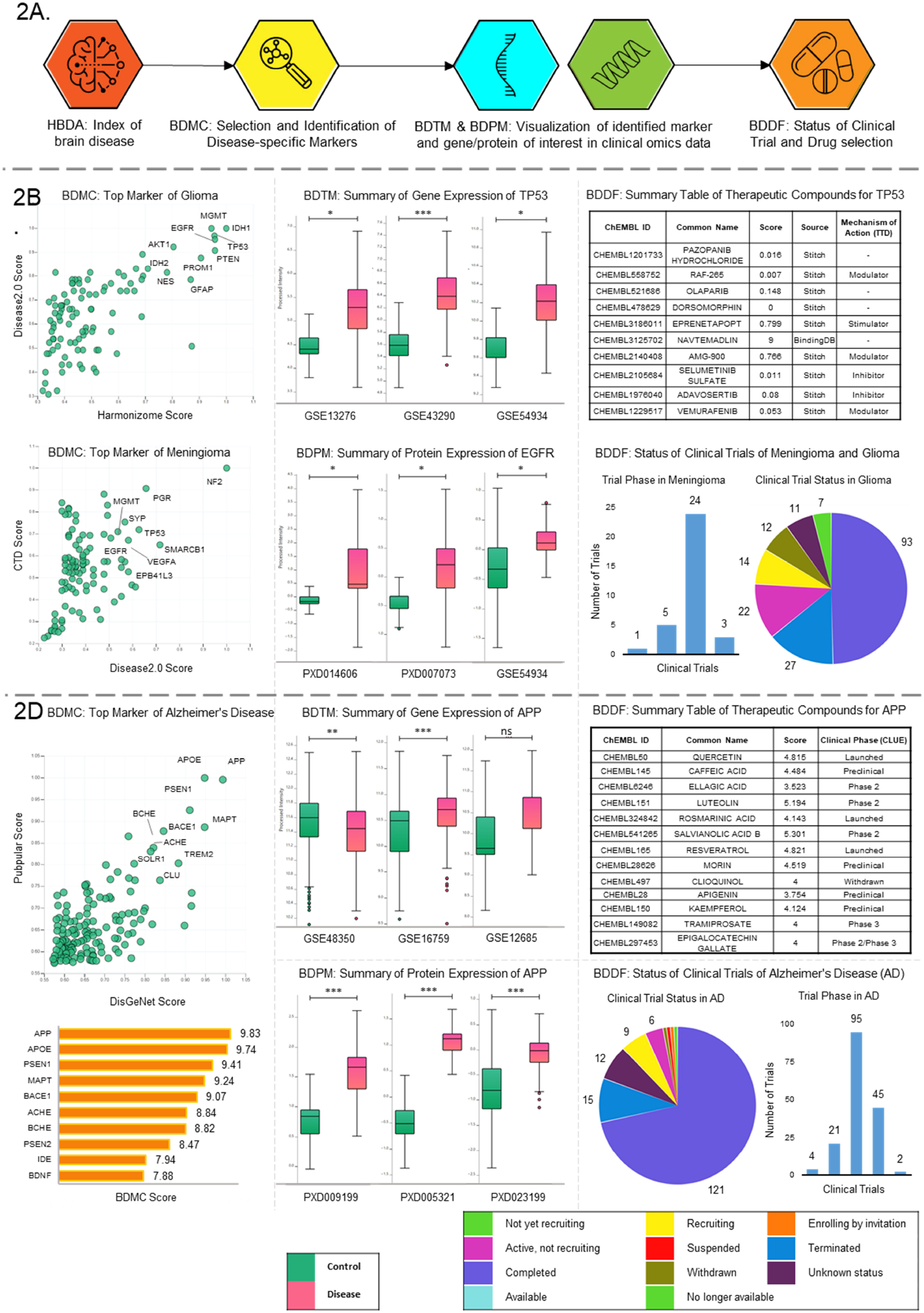

Amyloid Precursor Protein (APP) is a marker of Alzheimer’s disease (AD), a complex neurodegenerative disorder characterized by memory loss due to the deposition of the amyloid-beta peptide (42). APP has been found to be the top marker with a BDMC score of 9.825 for AD under BDMC. BDTM and BDPM integrates a total of 7 transcriptomics and 7 proteomics datasets. APP has showed upregulated in most of the datasets along with significant p-alue. The Clinical Trial section of BDDF recorded a total of 167 studies which showed one completed Phase 4 trial (NCT01397539), the investigation of BIIB037 (ADUCANUMAB). In addition, the total number of drug and chemical compounds against APP for AD are 1543 under the Disease Therapeutic Compounds section of BDDF (Figure 2C).

## DISCUSSION AND FUTURE DEVELOPMENTS

The development and integration of five new domains in BrainProt along with the modification of IBPM make it a comprehensive knowledge base with a robust, dynamic, and simple visualization platform for different Human Brain Diseases. BrainProt is freely available, open access database and developed with manually curated, analyzed, and reviewed data. This the first and only database that emphasizes marker identification and selection based on ranking by providing a list of 5 different scores and a combined BDMC Score for the 56 different brain diseases. In addition to this, BrainProt is the first knowledgebase that allows exploration of 52 omics (proteomics and transcriptomics) datasets mined, downloaded, and analyzed with a same analysis pipeline from 11 and 6 alarming diseases of the human brain under BDTM and BDPM sections respectively; however, all popular databases have been focused on and highlighted Omics Data of Glioma, Glioblastoma, or any one of the alarming brain diseases. The visualization of these two domains provides inter-disease and inter-dataset comparison along with easy access to more than 1,800 sample meta-data information for further clinical correlation. Lastly, BrainProt is the only knowledgebase that provides detailed information with visualization plots of more than 2,100 Clinical Trials with their phases and status information of more than 50 diseases along with exploration of Drug and Chemical compounds under BDDF.

Overall, BrainProt has been designed considering the usage of a vast community. The multiple intelligible plots, simple tables, user-friendly platform, hypertext links, rapid response, extensive Help/FAQ sections, and API-based data accessibility make BrainProt an exemplary platform for researchers worldwide. The objectives of BrainProt are to support and follow the footsteps of the HBPP (Human Brain Proteome Project) of the Human Proteome Organization (HUPO) to extend the magnitude and curtail the limits of knowledge of the Human Brain and its associated diseases.

## LIMITATIONS OF THE STUDY

The outcome and interpretation of the study have been solely dependent on the datasets and associated clinical information published and available in different repositories. New hypothesis, interpretation and markers generated from BrainProt should be validated before implementing or correlating for any kind of clinical practices. To the best of our knowledge, the BrainProt provides a foundation platform to assist and accelerate the neuroscience research,intending to expand the magnitude and push the limits of understanding of the Human Brain.

## ACKNOWLEDGMENT

We are grateful to the repositories and databases available which has been the major support to build the portal. We are also grateful to MASSFIITB, Proteomics Lab, IIT Bombay for their complete support during the development of the database.

